# Bulk flow of cerebrospinal fluid observed in periarterial spaces is not an artifact of injection

**DOI:** 10.1101/2020.11.09.374512

**Authors:** Aditya Raghunandan, Antonio Ladrón-de-Guevara, Jeffrey Tithof, Humberto Mestre, Maiken Nedergaard, John H. Thomas, Douglas H. Kelley

## Abstract

Cerebrospinal fluid (CSF) flowing through periarterial spaces is integral to the brain’s mechanism for clearing metabolic waste products. Experiments that track tracer particles injected into the cisterna magna of mouse brains have shown evidence of pulsatile CSF flow in pial periarterial spaces, with a bulk flow in the same direction as blood flow. However, the driving mechanism remains elusive. Several studies have suggested that the bulk flow might be an artifact, driven by the injection itself. Here, we address this hypothesis with new *in vivo* experiments where tracer particles are injected into the cisterna magna using a dual-syringe system, with simultaneous injection and withdrawal of equal amounts of fluid. This method produces no net increase in CSF volume and no significant increase in intracranial pressure. Yet, particle-tracking reveals flows in the pial periarterial spaces that are completely consistent with the flows observed in earlier experiments with single-syringe injection.

## Introduction

Cerebrospinal fluid (CSF) flowing in perivascular spaces (PVS) – annular tunnels that surround the brain’s vasculature — plays a crucial role in clearing metabolic waste products from the brain ***(Iliff et al., 2012; Xie et al., 2013**).* The failure to remove such waste products, including toxic protein species, has been implicated in the etiology of several neurological disorders, including Alzheimer’s disease *(**Iliff et al., 2012; Peng et al., 2016**).* Recently, *in vivo* experiments that combine two-photon microscopy and flow visualization in live mice have used the motion of fluorescent microspheres injected into the cisterna magna (CM) to measure the flow of CSF through the periarterial spaces surrounding pial (surface) arteries. The results show pulsatile flow, in lock-step synchrony with the cardiac cycle and with an average (bulk) flow in the same direction as that of the arterial blood flow ***Bedussi et al. (2017); Mestre et al. (2018b)**.* Characterizing the flow, however, is easier than determining its driver. Although arterial pulsation has long been considered as a possible driving mechanism for the bulk flow *(**Bilston et al., 2003; Hadaczek et al., 2006; Wang and Olbricht, 2011; Iliff et al., 2013; Thomas, 2019**),* that notion remains controversial *(**Diem et al., 2017; Kedarasetti et al., 2020a; van Veluw et al., 2020***), and other mechanisms are possible.

One such mechanism is the injection of tracers into the CM, which might cause a pressure gradient that drives a flow in the surface PVS *(**Smith et al., 2017; Smith and Verkman, 2018; Croci et al., 2019; Sharp et al., 2019; van Veluw et al., 2020; Vinje et al., 2020; Kedarasetti et al., 2020a**).* Injection of CSF tracers is known to raise the intracranial pressure (ICP) by 1 – 3 mmHg ***(Iliff et al., 2013; Mestre et al., 2020**),* consistent with the fact that a volume of fluid is being added to the rigid skull *(**Hladky and Barrand, 2018; Bakker et al., 2019**).* If that ICP increase is not uniform, the resulting pressure gradient could drive fluid into low-resistance pathways such as surface periarterial spaces *(**Faghih and Sharp, 2018; Bedussi et al., 2017**).* In that case, the bulk flows observed in detail by ***Mestre et al. (2018b)*** might have been artifacts of the injection. ***Mestre et al. (2018b)*** showed that the flows did not decay over time, as would be expected if they were injection artifacts, but given that injection artifacts have been suggested in several more recent publications, we decided to test the hypothesis with additional *in vivo* experiments, essentially identical to the earlier experiments *(**Mestre et al., 2018b**)*, but employing a new particle-injection method.

The new injection protocol, illustrated in Figure 1 b, employs a dual-syringe system to infuse the tracer particles. In this system, two cannulae connected to synchronized syringe pumps are inserted into the CM; one line injects fluid in which the tracer particles are suspended, while the other line simultaneously withdraws an identical amount of fluid at the same volumetric flow rate. Thus, no net volume of fluid is added to the intracranial compartment, and hence we expect no significant change in ICP. We use two-photon microscopy to visualize the motion of the fluorescent tracer particles and measure the flow in the surface periarterial spaces using particle tracking velocimetry. We also simultaneously measure changes to ICP while monitoring heart and respiration rates. We compare the flow characteristics measured under the new protocol with those measured previously using the traditional single-syringe injection *(**Bedussi et al., 2017; Mestre et al., 2018b**)* (depicted in Fig. 1 a). (For this comparison, the data from Mestre et al.(***Mestre et al., 2018b***) analyzed here are from the control mice, not the hypertension mice.) Our new results are completely consistent with the previous results. With the new infusion protocol, the flow is again pulsatile in nature, in step with the cardiac cycle, with a net (bulk) flow in the direction of arterial blood flow. We find nearly identical mean flow speeds and other flow characteristics with the new infusion protocol. Our new experiments confirm that the flows we observed in periarterial spaces in our earlier experiments are natural, not artifacts of the tracer infusion, and provide additional statistical information about these flows.

**Figure 1.**
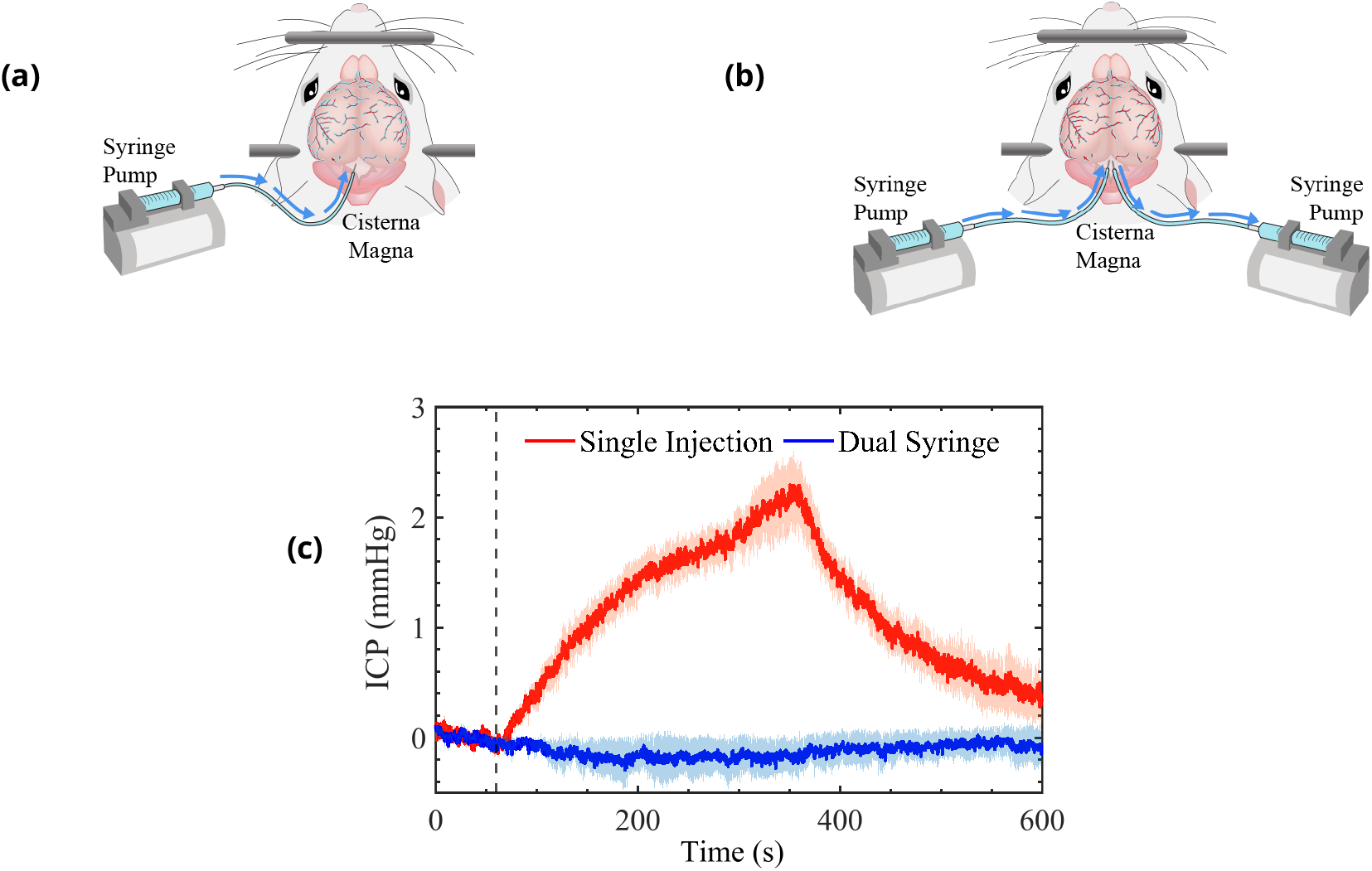
Schematic representation of the cisterna magna injection using (a) the single-syringe protocol for injection of 10 *μ*L at 2*μ*L/min and (b) the dual-syringe protocol for simultaneous injection and withdrawal of 20 *μ*L at 2*μ*L/min. The effect of single-injection and dual-syringe tracer infusion upon intracranial pressure (ICP) is shown in (c). The ICP was monitored continuously during injection of CSF tracers into the CM of mice. Injection begins at 60 s, indicated by the vertical dashed line. Single-injection infusion of 10 *μ*Lata rate of 2 *μ*L/min resulted in a mild change of – 2.5 mmHg in ICP, whereas little or no change in ICP was observed during the simultaneous injection and withdrawal in the dual-syringe protocol. Repeated measures two-way analysis of variance (ANOVA) was performed; interaction P value < 0.0001; n = 5 mice for single-injection and n = 6 mice for dual-syringe. The shaded regions above and below the plot lines indicate the standard error of the mean (SEM).

## Results

### Changes in intracranial pressure

In a group of mice, we evaluated the effect of tracer infusion upon ICP. A 30-gauge needle was inserted stereotactically into the right lateral ventricle and connected to a pressure transducer to monitor ICP during CSF tracer injection into the CM, using both the single-injection (n = 6 mice) and dual-syringe (n=5) protocols (Fig. 1c). In agreement with prior studies using similar single-injection protocols *(**Iliff et al., 2013; Xie et al., 2013; Mestre et al., 2020**),* we found that the injection of 10 *μ*L of CSF tracer into the CM at a rate of 2 *μ*L/min resulted in a mild elevation of ICP (~ 2.5 mmHg) that relaxed to baseline values within 5 min of the cessation of injection (Fig. 1 c). On the other hand, when ICP was measured during the dual-syringe infusion, we observed that the simultaneous injection of the tracer and withdrawal of CSF did not significantly alter ICP (Fig. 1 c), as expected given the absence of any net change in the volume of fluid in the intracranial CSF compartment. Based upon these findings, we conducted intracisternal infusion of fluorescent microspheres into the CM using the dual-syringe protocol to perform particle-tracking studies and determine the characteristics of CSF flow in the absence of any transient elevation of ICP caused by the infusion protocol.

### Flow measurements in perivascular spaces

We studied the motion of tracer particles infused with the new dual-syringe protocol (lower panels in Fig. 2) and compared it with the motion of tracer particles observed by *(**Mestre et al., 2018b**)* using the single-injection protocol (upper panels in Fig. 2), using particle tracking to examine flow of CSF in pial periarterial spaces.

**Figure 2.**
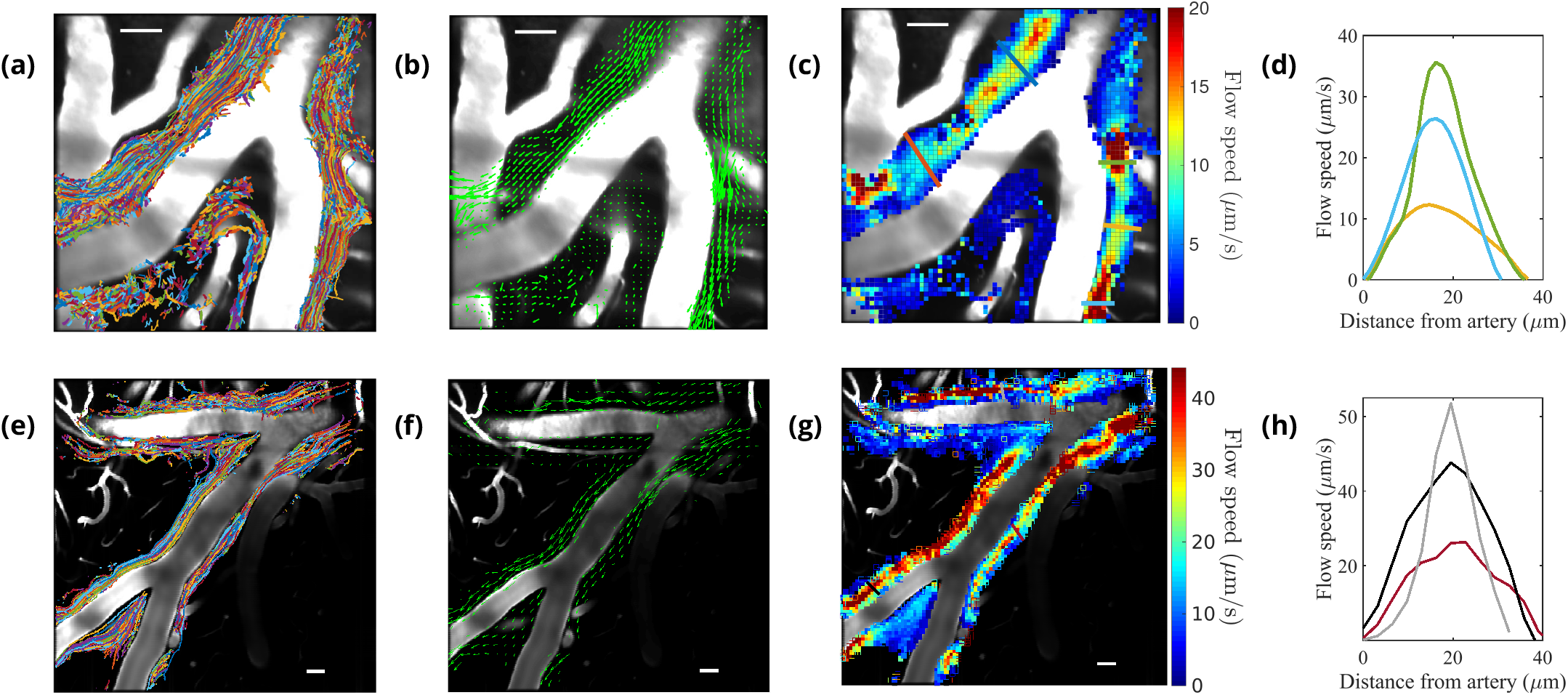
Particle tracking velocimetry in surface periarterial spaces using the single-injection method (panels in first row(***Mestre et al., 2018b***)) and the new dual-syringe method (second row). The superimposed particle tracks shown in panels (a) and (e) have similar, continuous spatial distributions and show similar sizes of the perivascular spaces. The time-averaged velocity fields shown in panels (b) and (f) both show net flow of fluid in the same direction as the blood flow. The flow-speed distributions plotted in panels (c) and (g) show comparable speeds, with the fastest flow at the center of the imaged periarterial space and the slowest flow near the boundaries. Panels (d) and (h) show average flow-speed profiles across the corresponding colored lines spanning the PVS in panels (c) and (g), smoothed by interpolation. The parabolic-like nature of these velocity profiles is what is expected for viscous flow in an open channel. Scale bars indicate 50 *μ*m.

The periarterial spaces of the cortical branches of the middle cerebral artery (MCA) were chosen for imaging. In the new protocol, particles appeared in the visualized spaces *~* 300 s after infusion was complete. This time scale is similar to that in our previous report *(**Mestre et al., 2018b**)* of 292 ± 26s, but particle counts were lower than those observed using the single-injection technique, likely because some of the injected particles were siphoned into the withdrawal line of the dualsyringe setup. However, a sufficient number of particles made their way into the PVSs to enable rigorous flow measurements (see Supplemental Video S1). Results obtained from the particle tracking analysis are shown in Figure 2. Each of the six experiments using the new protocol lasted at least 10 minutes and allowed us to track at least 6200 particles. An example of the superimposed particle tracks imaged in an experiment is shown in Figure 2e. The particle tracks are mostly confined to the perivascular spaces surrounding the artery, occasionally crossing from one side of the artery to the other. The distribution of particle tracks is spatially continuous across the width of the imaged PVSs under both infusion methods (Figs. 2a *(**Mestre et al., 2018b**)* & 2e), reaffirming that surface periarterial spaces are open, rather than porous, spaces *(**Min-Rivas et al., 2020**).* The direction of the observed fluid flow in the different branches is indicated by the arrows in Figures 2b and 2f. If injection were driving the flow, we would expect to observe dominant directional transport of tracer particles only when using the single-injection method, and little or no transport when using the dual-syringe method. The time-averaged (bulk) flow for both infusion methods is in the same direction as that of the blood flow, providing evidence that CSF flow in perivascular spaces is not caused by the injection. For both infusion methods, we observed no net flow in the direction opposite to that of blood flow, as some recent reports have suggested *(**Aldea et al., 2019; van Veluw et al., 2020)**.* Figure 2g shows that the average flow speed in pial (surface) periarterial spaces varies across the PVS, consistent with prior reports *(**Mestre et al., 2018b**)* shown in Figure 2c. The velocity prof le is parabolic-like (Figs. 2d and 2h);the flow is fastest (~50 ***μ***m/s) at the center of the PVS and slows to zero at the walls. This parabolic-like shape is consistent with laminar, viscous-dominated flow of CSF through an open annular space, and not through a porous medium, indicating that pial periarterial spaces are open ***Min-Rivas et al. (2020)**.*

Further analysis of the data obtained from particle tracking demonstrates the close similarity between the flows observed in the two protocols, as shown in Figure 3. A time-history of the measured flows — quantified by the spatial root-mean-square velocity computed at each instant of time *(**V_rms_***) — portrays very similar behavior over times much longer than the time it takes for the ICP to return to normal after the infusion (Fig. 3a). (The times shown here begin when particles were first seen or when the imaging was started: these times differ by less than 1 minute and so do not affect the results significantly.) The pulsatile nature of the flow at small time scales is depicted in Figures 3b and 3c. If injection-induced elevated ICP were driving the flow, we would observe large ***V_rms_*** values early in the single-injection experiments, followed by an exponential decay, and we would observe little or no flow in the dual-syringe experiments, in which the ICP remains unchanged. Since we observe very similar trends in the time-history profiles in both infusion protocols, the mechanisms driving the flow are apparently independent of the infusion method.

**Figure 3.**
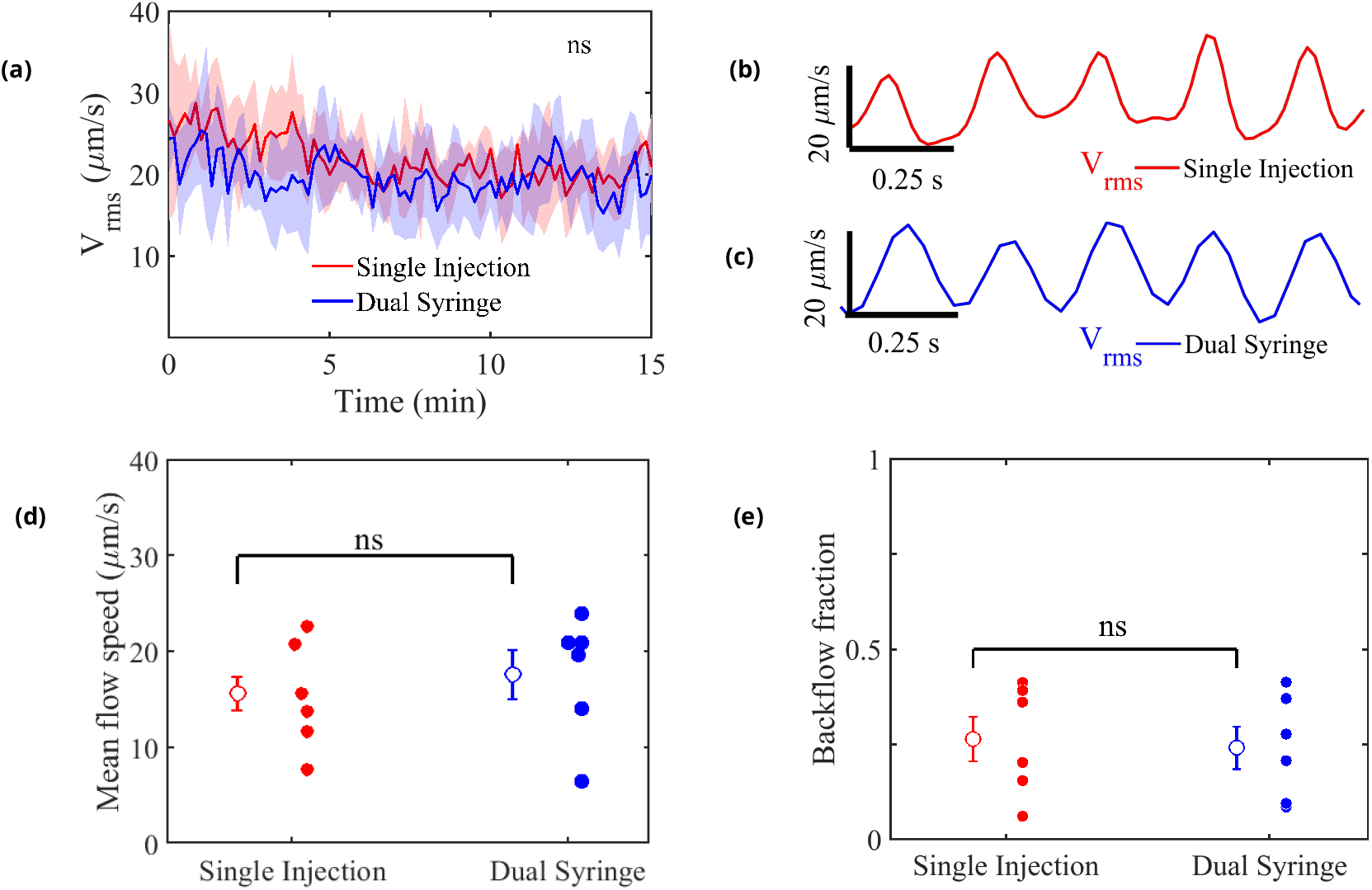
Measured flow characteristics. Panel (a) shows *V_**rms**_* over the course of the velocity measurements for both infusion methods. Repeated measures two-way ANOVA was performed;ns, not significant;n = 5 mice for single-injection and n = 6 mice for dual-syringe. The solid lines represent the mean value of *V_**rms**_* and the shaded area represents the standard error of the mean within each time bin. The pulsatility of typical measured flows is depicted in panels (b) and (c). Panel (d) shows mean downstream flow speeds and panel (e) shows backflow fractions for the individual experiments, with overall mean values shown as open circles (and bars showing the standard error of the mean). The nearly identical values for the two protocols demonstrate that the flow is independent of the injection method employed. Unpaired Student’s *t*-test was performed;n = 5 or 6 mice per group;ns, not significant;mean ± SEM.

Figure 3d shows mean flow speeds computed by averaging the downstream velocity component over space and time for each experiment. The overall mean flow speed (open circles) is 15.71 ± 6.2 *μ*m/s for all the single-injection experiments and 17.67 ± 4.42 *μ*m/s for all the dual-syringe experiments, values that differ by less than the standard error of the mean in either set of experiments. Significantly greater differences in mean flow speed are caused by animal-to-animal variations than by changing from single-injection to dual-syringe methods. These values are also nearly identical to the mean speed of 17 ± 2 *μ*m/s reported by ***Bedussi et al. (2017)***, from exper-iments that used a single-injection protocol with a lower injection rate. The mean flow speeds represent the speeds at which tracer particles (or cerebrospinal fluid) are transported in the direction of arterial blood flow (downstream), and presumably into the brain. If the observed flows were injection-induced, we would expect faster mean flows with the single-injection method than with the dual-syringe method.

We also computed a ‘backflow fraction’ for each experiment, as the fraction of the downstream velocity measurements showing motion in the retrograde direction (opposite that of the blood flow): the results are shown in Figure 3e. An injection-driven flow would be dominantly unidirectional and would exhibit a much smaller backflow fraction. However, the backflow fraction is nearly identical: 0.26 ± 0.059 for single-injection and 0.24 ± 0.056 for dual-syringe infusion respectively. As with flow speed, mean values differ by less than the standard error of the mean, so animal-to-animal variations exceed the effects of changing the injection protocol. The nearly identical mean flow speeds and backflow fractions further demonstrate that the observed flows are natural, and not artifacts of the infusion.

### Pulsatile flow is regulated by the cardiac cycle

It has been variously suggested that CSF flow might be driven by the cardiac cycle, the respiratory cycle, or perhaps both *(**Rennels et al., 1985; Hadaczek et al., 2006; Yamada et al., 2013; Bedussi et al., 2017)***, with evidence indicating much stronger correlation with the cardiac cycle ***(Iliff et al., 2013; Mestre et al., 2018b**).* We used the simultaneous measurements of the electrocardiogram (ECG) and respiration in conjunction with particle tracking to determine the relative importance of the cardiac and respiratory cycles *(**Santisakultarm et al., 2012**),* and also to see if there is any difference in these relationships between the two infusion methods (Fig. 4).

**Figure 4.**
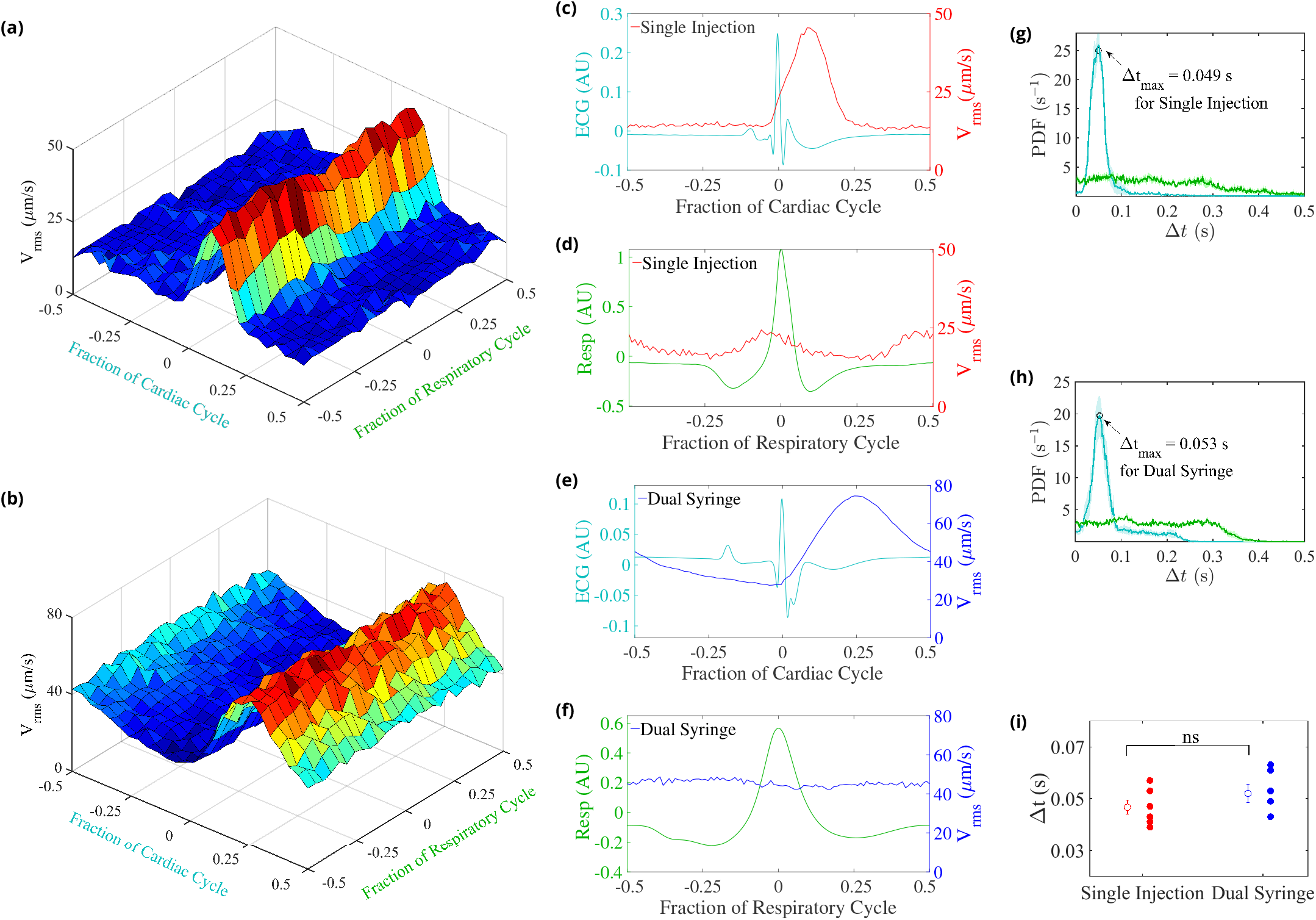
CSF velocity variations over the cardiac and respiratory cycles. Panels (a) and (b) show the measured ***V_rms_*** conditionally averaged over the cardiac and respiratory cycles, based on the synchronized measurements of ECG, respiration, and velocity, for the single-injection (a) and the dual-syringe (b) protocols. Panel (c) for single injection and panel (e) for dual syringe both show that the peaks in the ECG are immediately followed by peaks in ***v_rms_***, indicating a strong correlation between heart rate and fluid motion in both injection protocols. No consistent trends are seen when ***v_rms_*** is averaged over the respiratory cycle, as shown in panels (d) and (f)· Panels (g) and (h) show the mean and the standard error of the mean of probability density functions of the delay time Δt between the peak in the cardiac (cyan) or respiration (green) cycle and the subsequent peak in ***v_rms_***, for single-injection (*n*=5) and dual-syringe (*n* = 6) methods respectively. Panel (i) shows the average Δt between peaks in the cardiac cycle and *v_rmj_* for both protocols; in both, the peak in *v_rms_* typically occurs ~ 0.05 s after the peak in the cardiac cycle. Unpaired Student’s t-test was performed; n = 5 or 6 mice per group; ns, not significant; mean ± SEM.

We find that the measured time-dependent components of flow quantities such as ***V_rms_*** are strongly modulated by the cardiac cycle but only weakly by respiration (Figs. 4a ***(Mestre et al., 2018b)*** & 4b). This strong correlation of the pulsatile component of flow with the heart rate is exhibited under both the single-syringe and dual-syringe protocols, as shown in Figures 4c and 4e, where the peak in the ***V_rms_*** occurs soon after the peak in the cardiac cycle. Probability density functions of Δt, the delay time between peaks in ***V_rms_*** and cardiac/respiratory cycles, also predict a much greater likelihood of peaks in ***V_rms_*** following the peak in the cardiac cycle (Figs. 4g & 4h). We observe nearly identical average delay times of ~ 0.05 s between peaks in ***V_rms_*** and the cardiac cycle for both protocols (Fig. 4i). No such correlation is observed when the ***V_rms_*** is conditionally averaged over respiration cycles (Figs. 4d and 4f). These observations corroborate prior reports that the cardiac cycle drives the dominant oscillatory component of CSF flow in the surface periarterial spaces, unaffected by injection protocol.

## Discussion

Healthy removal of metabolic waste from the brain is believed to occur via circulation of CSF, which enters brain tissue through perivascular spaces surrounding pial arteries ***(Rasmussen et al., 2018; Reeves et al., 2020; Nedergaard and Goldman, 2020**).* Whereas experiments in live mice have shown that fluid is pumped in the direction of blood flow and into brain, perhaps by forces linked to the pulsation of arterial walls, several published papers have hypothesized that the observed flows might instead be artifacts of non-natural elevation of ICP caused by tracer infusion into the cisterna magna. In this study, we designed a new infusion protocol that enabled tracer-particle infusion with no net addition of fluid and near-zero changes in ICP. We used two-photon microscopy and particle-tracking velocimetry, and found flows of CSF in the surface periarterial spaces that are statistically identical to the flows found earlier using the single-injection protocol. The measured flows are pulsatile, viscous-dominated, laminar flows, with mean flow in the direction of blood flow in the cerebral arteries. Our flow visualization techniques and synchronized measurements of ICP and heart and respiration rates enabled us to show that the observed flows are not driven by pressure differences induced by the tracer infusion.

Our new experiments provide several lines of evidence that the observed bulk flow is not induced by tracer infusion methods currently used. Tracer infusion at rates of 1 – 2 *μ*L/min have typically been employed *x(**Bedussi et al., 2017; Mestre et al., 2018b**)* to add 10 *μ*L of fluid to the subarachnoid space. Although this addition of fluid is greater than the natural CSF production rate (0.38 *μ*l/min *(**Oshio et al., 2005***)), and it induces a small increase in ICP, our results show that the ICP returns to its baseline value within 5 minutes after injection is completed. If infusion were propelling the mean flow, particle transport would cease after the ICP reverted to normal, or would not occur at all with dual-syringe injection, where the ICP is not affected. However, we typically observe particles being transported along the periarterial spaces for 30 min, long after the return of ICP to its baseline value in the single-syringe experiments and at comparable times in the dual-syringe experiments.

Elevated ICP levels create large pressure differences across the brain, but these pressure differences will undergo exponential decay because of the brain’s compliance and proclivity to achieve stasis. If this exponential relaxation of ICP were to drive fluid flow, the measurements from particle tracking would reflect this decay, exhibiting fast flows at early times which then gradually subside. However, our measurements show that the mean flow remains nearly constant and similar over periods that are 2 to 3 times longer than the infusion time, for several healthy mice and both infusion protocols. Variability is probably due to physiological differences between different mice. Moreover, if infusion-driven flows were prevalent, we would expect to see much faster mean flow speeds. Yet, with the new dual-syringe method, with no net infusion, we observe flow speeds that are nearly identical to those observed in our earlier study *(**Mestre et al., 2018b**),* and very close to those in a study that used injection rates many orders of magnitude smaller *(**Bedussi et al., 2017**).* An infusion-driven flow would also be unidirectional, with no retrograde motion, but we observed consistent backflows of particles for both infusion protocols. Finally, if the ICP elevation induced by the single-injection protocol were responsible for the tracer penetration into the brain, then variations associated with arousal state *(**Xie et al., 2013**),* anesthesia *(**Hablitz et al., 2019**),* blood pressure ***Mestre et al. (2018a)***, and other biological mechanisms would not have occurred.

Our results confirm that the cardiac cycle — not respiration — drives the oscillatory component of the observed periarterial flows. The peaks of ***V_rms_*** that we measured across specimens, for both infusion protocols, appear shortly after the peaks in the cardiac cycle, but are not correlated with the respiratory cycle. Probability density functions show that the delay times between the peaks in the cardiac cycle and the peaks in ***V_rms_*** are nearly identical for the two infusion methods. Although we present compelling evidence that the cardiac cycle drives the purely oscillatory component of the pulsatile flow, we cannot rule out other natural mechanisms that might be driving the average (bulk) flow, such as CSF production, functional hyperemia *(**Kedarasetti et al., 2020b**),* or vasomotion (***van Veluw et al., 2020; Kiviniemi et al., 2016***). We do conclude, however, that the currently employed methods of tracer infusion are not responsible for the observed flows.

## Materials and Methods

### Animals and surgical preparation

All experiments were approved and conducted in accordance with the relevant guidelines and regulations stipulated by the University Committee on Animal Resources of the University of Rochester Medical Center (Protocol No. 2011-023), certified by Association for Assessment and Accreditation of Laboratory Animal Care. An effort was made to minimize the number of animals used. We used 8- to 12-week-old male C57BL/6 mice acquired from Charles River Laboratories (Wilmington, MA, USA). In all experiments, animals were anesthetized with a combination of ketamine (100 mg/kg) and xylazine (10 mg/kg) administered intraperitoneally. Depth of anesthesia was determined by the pedal reflex test. Once reflexes had ceased, anesthetized mice were fixed in a stereotaxic frame for the surgical procedure and body temperature was kept at 37°C with a temperature-controlled warming pad.

### Dual-syringe protocol

For *in vivo* imaging, anesthetized mice were fixed in a stereotaxic frame and body temperature was maintained at 37.5°C with a rectal probe-controlled heated platform (Harvard Apparatus). Two 30-gauge needles were inserted into the cisterna magna, as previously described ***(Xavier et al., 2018)**.* Briefly, the dura mater of mice was exposed after blunt dissection of the neck muscles so that a cannula could be implanted into the cisterna magna (CM), which is continuous with the subarachnoid space. A cranial window was prepared over the right middle cerebral artery (MCA) distribution. The dura was left intact, and the craniotomy (≃4 mm in diameter) was filled with aCSF, covered with a modified glass coverslip, and sealed with dental acrylic. Afterwards, two 30-gauge needles were inserted into the cisterna magna, as described above. Using a syringe pump (Harvard Apparatus Pump 11 Elite), red fluorescent polystyrene microspheres (FluoSpheres^T^*M* 1.0 *μ*m, 580/605 nm, 0.25% solids in aCSF, Invitrogen) were infused up to a total volume of 20 *μ*L via one of the cisterna magna cannulae while CSF was simultaneously withdrawn through the other cannula at an equal rate of 2 *μ*L/min with a coupled syringe pump.

### Intracranial pressure measurements

Anesthetized mice were fixed in a stereotaxic frame, and two 30-gauge needles were inserted into the cisterna magna, as described above. A third cannula was inserted via a small burr hole into the right lateral ventricle (0.85 mm lateral, 2.10 mm ventral and 0.22 mm caudal to bregma). Mice were then placed in a prone position. In the first set of experiments, 10 ***μ***L of artificial CSF (aCSF) was injected into the CM at a rate of 2 ***μ***L/min via one of the CM cannulae using a syringe pump (Harvard Apparatus Pump 11 Elite). In the second set of experiments, aCSF was injected at the same rate while withdrawing CSF from the cisterna magna via the other CM cannula at an equal rate using a coupled syringe pump (Harvard Apparatus Pump 11 Elite). In both experiments, intracranial pressure (ICP) was monitored via the ventricle cannulation connected to a transducer and a pressure monitor (BP-1, World Precision Instruments). ICP was acquired at 1 kHz, digitized, and monitored continuously for the duration of the infusion experiments with a DigiData 1550B digitizer and AxoScope software (Axon Instruments).

### *In vivo* two-photon laser-scanning microscopy

Two-photon imaging was performed using a resonant scanner B scope (Thorlabs) with a Chameleon Ultra II laser (Coherent) and a 20× water immersion objective (1.0 NA, Olympus). Intravascular FITC-dextran and red microspheres were excited at a 820 nm wavelength and images were acquired at 30 Hz (ThorSync software) simultaneously with physiological recordings (3 kHz, ThorSync software), as previously described *(**Mestre et al., 2018b**).* To visualize the vasculature, fluorescein isothiocyanate-dextran (FITC-dextran, 2,000 kDa) was injected intravenously via the femoral vein immediately before imaging. Segments of the middle cerebral artery were distinguished on the basis of morphology: surface arteries passing superficially to surface veins and exhibiting less branching at superficial cortical depths. ECG and respiratory rate were acquired at 1 kHz and 250 Hz, respectively, using a small-animal physiological monitoring device (Harvard Apparatus). The signals were digitized and recorded with a DigiData 1550A digitizer and AxoScope software (Axon Instruments).

### Image processing

Images with spatial dimensions 512 x 512 were obtained from two-photon microscopy. Each image is 16-bit with two channels, red and green. The FITC-dextran injected in the vasculature is captured via the green channel while the red channel is used to image the fluorescent microspheres flowing in the perivascular spaces. Image registration via rigid translation is performed on each image in the time series to account for movement by the mouse in the background. The image registration is implemented using an efficient algorithm in Matlab *(**Guizar-Sicairos et al., 2008**)* to an accuracy of 0.2 pixels. Erroneous correlations in the translation are manually corrected by linear interpolation. The translations obtained are sequentially applied to images that are padded with zero-value pixels. This ensures spatial dimension homogeneity across all images without modifying the image resolution. Particles are then detected by applying a minimum intensity threshold to each image. Typically, particles were resolved across 3-4 pixels in the image with spatial resolution of 1.29 ***μ***m.

### Particle-tracking velocimetry

The particles detected in each image were tracked using an automated PTV routine in MATLAB ***(Kelley and Ouellette, 2011; Ouellette et al., 2006**).* Briefly, the algorithm locates each particle with a sub-pixel accuracy and obtains a series of particle locations (particle tracks) for the entire duration of the recorded video. Particle velocities were calculated by convolution with a Gaussian smoothing and differentiation kernel. Stagnant particles that have adhered to the wall of the artery or the outer wall of the PVS, and hence no longer track the CSF flow, were masked in each image by subtracting a dynamic background image. This image was different for each frame and was computed by taking the average of 100 frames before and 100 frames after the given image. This method of masking was applied only to the dual-syringe data; the single-syringe data used a simpler masking approach with a single background image *(**Mestre et al., 2018b**).* Time-averaged flow velocities were obtained by segregating the imaged domain into a 70 x 70 grid, with a resolution of 7.5 x 7.5 pixels in each direction. All velocity measurements for a chosen time interval were binned based on their grid position. Average flow speeds were computed using bins with at least 15 measurements. The downstream velocity component was calculated as the dot product ***u · ŭ_avg_***, where ***u*** is the instantaneous particle velocity and ***ŭ_avg_*** is the field of unit vectors computed from the time-averaged flow field, in the direction of arterial blood flow.

### Statistical analysis

All statistical analyses were performed on GraphPad Prism 8 (GraphPad Software). Data in all graphs are plotted as mean ± standard error of the mean (SEM) over the individual data points and lines from each mouse. Statistical tests were selected after evaluating normality (D’Agostino Pearson omnibus test). When the sample size did not allow for normality testing, both parametric and nonparametric tests were performed and, in all cases, yielded the same result. Sphericity was not assumed; in all repeated measures, two-way ANOVAs and a Geisser-Greenhouse correction were performed. All hypothesis testing was two-tailed and exact P values were calculated at a 0.05 level of significance and stated in the figure legends.

## Acknowledgments

We thank Keith Sharp for recommending something like our dual-syringe experiments to us in July 2019. We also thank Dan Xue for drafting the schematic.

